# Multiple PDZ Domain Protein Maintains Patterning of the Apical Cytoskeleton in Sensory Hair Cells

**DOI:** 10.1101/2021.02.20.432099

**Authors:** Amandine Jarysta, Basile Tarchini

## Abstract

Sound transduction occurs in the hair bundle, the apical compartment of sensory hair cells in the inner ear. The hair bundle is formed of stereocilia aligned in rows of graded heights. It was previously shown that the GNAI-GPSM2 complex is part of a developmental blueprint that defines the polarized organization of the apical cytoskeleton in hair cells, including stereocilia distribution and elongation. Here we report a novel and critical role for Multiple PDZ domain (MPDZ) protein during apical hair cell morphogenesis. We show that MPDZ is enriched at the hair cell apical membrane, and required there to maintain the proper segregation of apical blueprints proteins, including GNAI-GPSM2. Loss of the blueprint coincides with misaligned stereocilia in *Mpdz* mutants, and results in permanently misshapen hair bundles. Graded molecular and structural defects along the cochlea can explain the profile of hearing loss in *Mpdz* mutants, where deficits are most severe at high frequencies.

## INTRODUCTION

Auditory function relies on the ability of sensory hair cells (HCs) in the auditory epithelium to detect sound vibrations. This is the role of the hair bundle, a highly organized brush of actin-based membrane protrusions, or stereocilia, located at the HC apical surface. Mutations affecting proteins involved in the morphogenesis or the maintenance of the hair bundle lead to hearing loss [1–4].

We and others showed that inhibitory Guanine nucleotide-binding proteins (G proteins) of the alpha family (GNAI1-3, collectively GNAI) bind the scaffold protein G Protein Signaling Modulator 2 (GPSM2) to form a complex essential for the polarized morphogenesis of the apical cytoskeleton in HCs [5–11]. Post-mitotic HCs are initially covered uniformly with microvilli, but planar polarized enrichment of GNAI-GPSM2 and Inscuteable (INSC) at the apical membrane defines a lateral/abneural region without protrusions, the bare zone [8]. GNAI-GPSM2 generates and expands the bare zone by antagonizing the polarity kinase aPKC that coincides with the microvillar HC surface. Complementary GNAI-GPSM2 vs aPKC distribution thus forms an apical blueprint that defines the lateral edge of the emerging hair bundle when microvilli grow into stereocilia [8]. As HCs mature, the GNAI-GPSM2 complex is then trafficked to the tips of stereocilia adjacent to the bare zone by the MYO15A motor [7, 9, 10]. At stereocilia tips, GNAI-GPSM2 promotes enrichment of MYO15A and other partners, conferring the first stereocilia row with its unique tallest identity [10]. Loss of GNAI or GPSM2 in mouse leads to stereocilia stunting, and the resulting immature hair bundle morphology in adults coincides with profound deafness [5, 7, 9]. In humans, mutations in *GPSM2* are responsible for the Chudley-McCullough syndrome where hearing loss is the principal affliction [12, 13].

GNAI-GPSM2 is enriched and polarized independently from planar cell polarity (PCP) proteins [6, 8], the machinery that regulates HC orientation via intercellular communication at apical cell-cell junctions [14, 15]. Accordingly, polarization of a subset of PCP proteins was recently proposed to depend on Wnt secretion, but severely disrupted cochlear elongation and morphogenesis in *Wntless* (*Wls*) mutants left GNAI-GPSM2 enrichment intact at the HC surface [16, 17]. Similarly, polarized segregation of GNAI-GPSM2 and aPKC was preserved in mutants lacking other polarity proteins that occupy the lateral apical HC junction: DAPLE/CCDC88C [18] and PARD3 [19]. It thus remains unclear how GNAI-GPSM2 is first polarized and then maintained at the HC apical membrane. Here, we hypothesized that GNAI-GPSM2 might more specifically interact with polarity proteins that occupy the HC apical membrane. To find possible candidates, we investigated proteins well-known to establish apico-basal polarization in other epithelial systems.

Multiple PDZ domain (MPDZ, or MUPP1) is a large adapter protein with 13 PDZ (PSD95/DLG1/ZO1) domains that mediates protein-protein interactions and was first reported as interacting with the HTR2C serotonin receptor [20, 21]. MPDZ is a scaffold protein that clusters binding partners in defined cell compartments, for example SynGAP and CaMKII at synapses [22], as well as Delta Like protein 1 and 4 (DLL1/4) and Nectin-2 at epithelial apical junctions [23]. MPDZ is a paralog of INADL/PATJ, a scaffolding protein that binds MPP5/PALS1, which in turn binds the integral protein CRB3 to form the Crumbs complex [24–27]. The Crumbs complex plays a highly conserved role in the establishment of epithelial apico-basal polarity [28, 29], including the formation of tight junctions in mammals [26, 27, 30–33]. MPDZ itself was reported as a component of tight junctions, and MPDZ can directly bind Claudins [34–36] as well as many of the INADL partners, including JAM1 [35, 37] and ZO1 [37]. Interestingly, like INADL, MPDZ can bind MPP5 to form an alternative Crumbs complex (CRB3-MPP5-MPDZ) with distinct cellular functions from the better-described CRB3-MPP5-INADL complex [27, 37]. Recently, MPDZ was shown to bind and cooperate with DAPLE at apical junctions for apical cell constriction in cell culture and during neurulation [38, 39].

In human and mice, mutations in *MPDZ* lead to multiple ependymal malformations that compromise the lining of the brain ventricles [40, 41]. These defects were initially proposed to result from defective ependymal tight junctions, but a recent study also uncovered a more specific role for MPDZ in the choroid plexus that secretes cerebrospinal fluid into the ventricles [42]. Interestingly, although barrier integrity was defective in both organs, MPDZ was only reported as apical in choroid plexus epithelial cells, and not visibly enriched at tight junctions [42]. Likely due to ependymal and/or choroid plexus dysfunction, *MPDZ* is one of the few genes linked to congenital hydrocephalus in humans [40, 43–45] and mice [41, 42]. Of note, a young patient carrying two distinct *MPDZ* mutations in *trans* also presented with sensorineural hearing loss, suggesting a potential role for MPDZ in the inner ear [45].

In this study, we identify a new function for MPDZ during HCs development. Surprisingly, MPDZ is not detected at apical cell junctions in the auditory epithelium, but at the HC apical membrane where it co-localizes with MPP5 and CRB3. We show that MPDZ is necessary to maintain the apical segregation of GNAI-GPSM2 and aPKC. Loss of this apical blueprint in *Mpdz* mutants coincides with misaligned stereocilia and dysmorphic hair bundles. Molecular and morphological defects persist in young adults and are likely responsible for severe hearing loss in *Mpdz* mutants.

## RESULTS

### MPDZ occupies the hair cell apical membrane with a lateral bias

To investigate a potential role for MPDZ in the auditory epithelium, we started by immunolabeling MPDZ at different developmental stages with a validated antibody targeting the third PDZ domain (PDZ3) [34]. As early as embryonic day (E) 17.5, MPDZ signal was obvious at the HC apex, spanning most of the apical membrane (Fig. 1A). MPDZ was most highly enriched at the bare zone, the flat region deprived of protrusions lying lateral to the tallest stereocilia [8]. MPDZ signal was lower on the medial side of the hair bundle, and absent or barely detectable at the position of the hair bundle itself. A similar MPDZ enrichment pattern was observed in neonate HCs at postnatal day (P) 0 and P4 (Fig. 1A). Of note, INSC, GNAI and GPSM2 are by contrast strictly enriched at the bare zone and are thus more obviously planar polarized than MPDZ [6, 8, 11].

**Figure 1.**
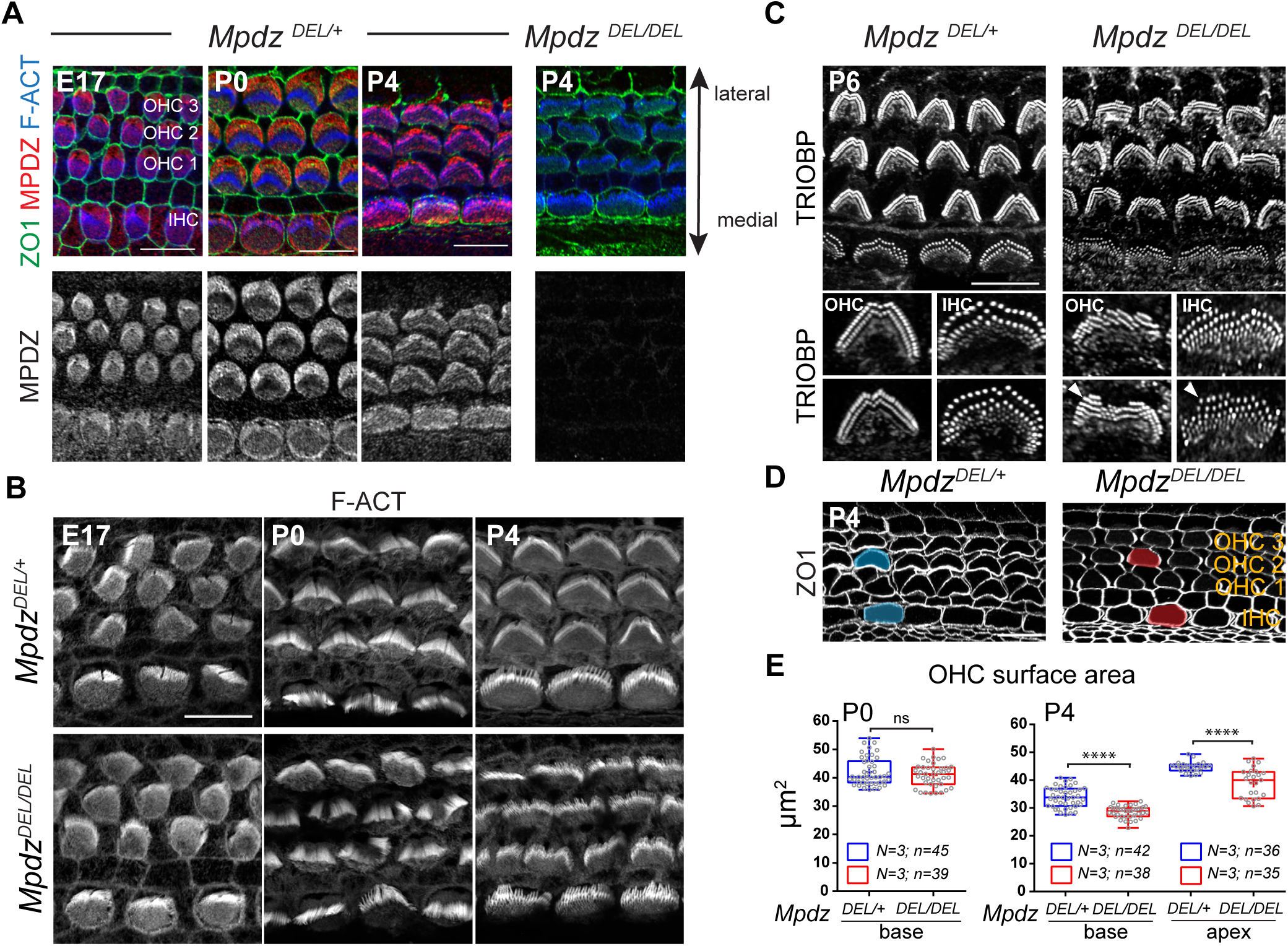
MPDZ occupies the hair cell apical membrane and is required for normal junction circumferences and stereocilia distribution in neonates. **A)** Flat mount views of the mouse auditory epithelium immunolabeled with MPDZ (red), ZO1 (green, apical junctions) and phalloidin (blue, F-actin) at E17, P0 and P4 (cochlear base). MPDZ signal is entirely lost in *Mpdz* mutants (*Mpdz^DEL/DEL^*). **B)** Phalloidin (F-actin) labeling of the hair bundle level (cochlear base; maximal projection). In absence of MPDZ, OHC and IHC hair bundles are misshapen from P0. **C)** TRIOBP labeling of the stereocilia rootlets at P6 (maximal projection). Stereocilia are misaligned and form interrupted or supernumerary rows in *Mpdz* mutants. **D)** ZO1 labeling of the apical junctions (cochlear base) at P4. In absence of MPDZ, the HC circumference is abnormally rounded, and the apical surface appears reduced. **E)** Apical OHC surface area at P0 and P4 based on ZO1 labeling. Here and in the following figures, graphical representations show a boxplot with 25-75% percentiles where external bars show the minimum and maximum, the internal bar shows the median and the cross shows the mean. All points are also shown. N = number of animals, n = total number of OHCs (Mann-Whitney U test; P0: p=0.4417 ^ns^; P4 base: p<0.0001 ****; P4 apex: p<0.0001****; ns, not significant). Scale bars are 10μm.

### Loss of MPDZ results in apical defects in developing hair cells

We obtained a constitutive *Mpdz* mouse mutant from the IMPC repository (C57BL/6N-*Mpdz^em1(IMPC)J^*/Mmucd, hereafter *Mpdz^DEL^*). This strain carries an indel in *Mpdz* exon 6 which is predicted to cause a frameshift resulting in an early truncation of the protein after the first of 13 PDZ domains. We did not observe lethality in *Mpdz^DEL/DEL^* animals, whereas previous *Mpdz* mutants died around 3-4 week of age (see Supplementary Table 1) [41, 42, 46]. Nevertheless, as expected, MPDZ antibody signal was entirely missing in the *Mpdz^DEL^*^/DEL^ auditory epithelium, establishing this strain as a proper loss-of-function model in HCs (Fig. 1A).

Based on its localization, we asked whether MPDZ could be involved in apical HC morphogenesis. We first imaged the hair bundle at embryonic and postnatal stages using conjugated phalloidin to reveal F-actin. At E17.5, stereocilia organization appeared comparable between *Mpdz* mutants and littermate controls at the cochlear base (Fig. 1B). By birth (P0) however and at P4, apical HC defects were obvious in outer HCs (OHCs), where hair bundles lacked their characteristic V-shape and were frequently split (Fig. 1B). Because 3-dimensional organization of protruding stereocilia can be confusing in flat-mounts, we labeled stereocilia rootlets at P6 with an antibody against TRIOBP [47]. We found that stereocilia were severely misaligned in *Mpdz* mutants (Fig. 1C). Stereocilia also formed incomplete supernumerary rows most obvious on the lateral side of the bundle at P6 (Fig. 1C, arrowhead). Inner HCs (IHCs) showed similar, but milder defects from P0 (Fig. 1B-C). Together, these results suggest that MPDZ acts at the apical membrane outside of the forming hair bundle and is important for the proper localization and cohesive, arrayed organization of stereocilia in postnatal HCs.

Labeling of apical junctions with ZO1 revealed an abnormally round and possibly reduced OHC circumference in P4 *Mpdz* mutants compared to control littermates (Fig. 1D). IHCs were not visibly affected. We quantified the apical surface area in OHCs and observed a significant reduction in *Mpdz* mutants at the cochlear base and apex at P4, but not at P0 (Fig. 1E). The OHC apical surface is known to decrease after P0 along with a remodeling of the cell circumference that closely mirrors hair bundle development [48]. MPDZ might thus regulate this process at the flat apical membrane, impacting both the placement of stereocilia and the shape of the HC junctions. GNAI and GPSM2 share MPDZ localization at the bare zone, and are also essential to define the lateral outline of the forming hair bundle [6, 8, 11], as well as the asymmetry of height and identities across stereocilia rows [7, 9, 10]. Next, we thus analyzed how GNAI-GPSM2 localized in *Mpdz* mutants.

### Loss of MPDZ causes a progressive delocalization of GNAI-GPSM2 at the hair cell surface

We assessed GNAI localization in HCs in the first postnatal week. At P2, GNAI was distributed normally in *Mpdz* mutant HCs along the whole cochlea (Fig. 2A). In contrast, at P4 many OHCs at the cochlear base showed medial delocalization of GNAI, whereas HCs at mid and apex cochlear positions retained normal GNAI distribution. At P7, all OHCs at the base and many OHCs at the mid position showed delocalized GNAI, and HCs at the apex retained normal GNAI distribution. We conclude that loss of MPDZ progressively alters apical GNAI distribution as HC differentiation progresses in time and spatially along a cochlear apex to base gradient [49]. Quantification of GNAI signal intensity in P4 OHCs confirmed a drastic delocalization from the lateral to the medial side of the hair bundle. Compared to control littermates, GNAI enrichment at the lateral bare zone decreased, and medial GNAI enrichment increased in *Mpdz* mutants (Fig. 2B).

**Figure 2.**
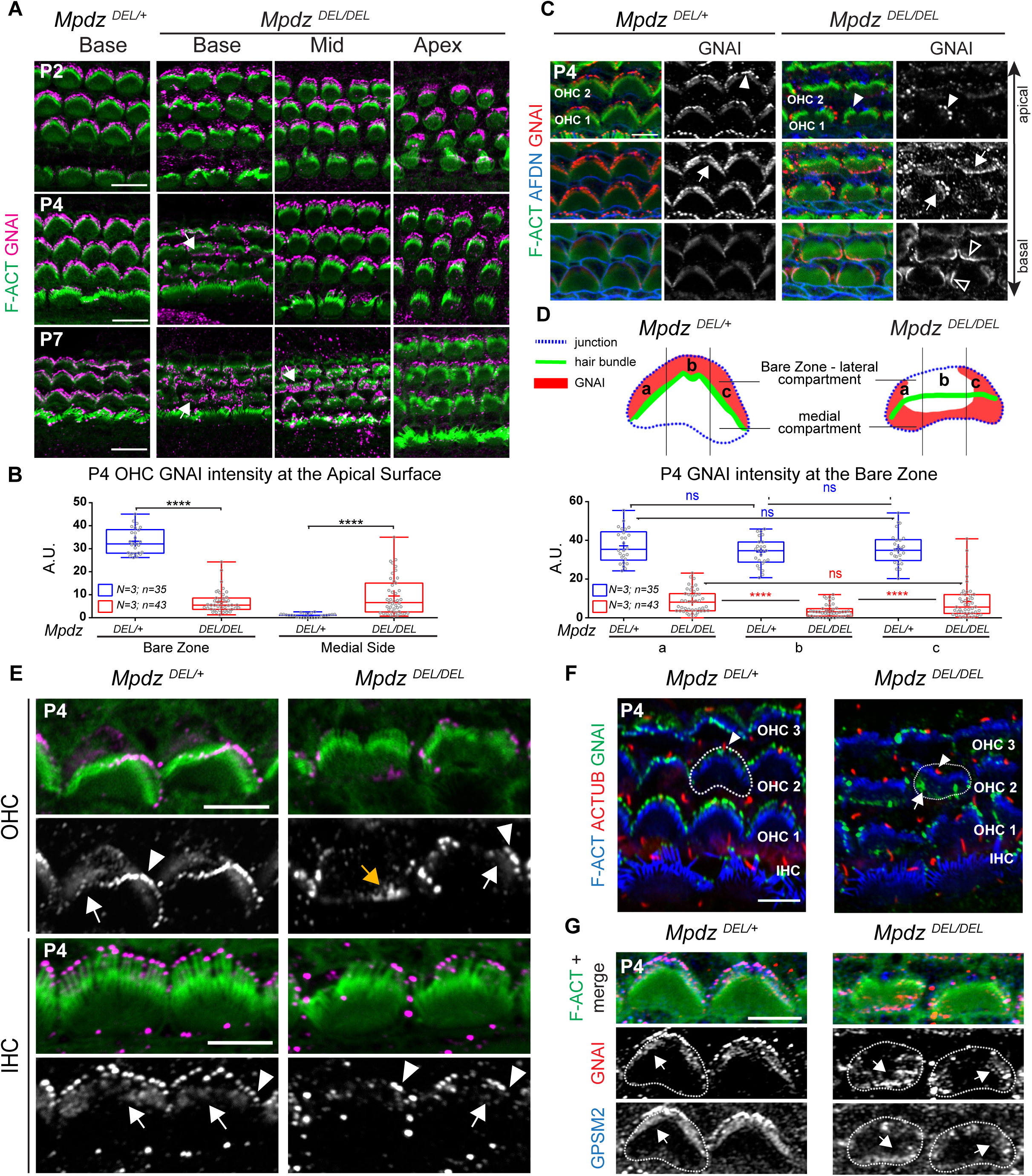
Apical enrichment of GNAI and GPSM2 is progressively reduced and delocalized in absence of MPDZ. **A)** GNAI immunolabeling (magenta) at different postnatal stages and positions along the cochlea (maximal projection). In *Mpdz* mutants, GNAI enrichment at the lateral bare zone and in neighboring stereocilia is compromised at the cochlear base at P4 (arrows), and this defect extends to the cochlear mid by P7. **B)** GNAI enrichment at the lateral (bare zone) and medial apical surface in P4 OHCs. N = number of animals, n = total number of OHCs (Mann-Whitney U test; Bare zone and medial side: p<0.0001 ****). **C)** GNAI labeling at P4 at 3 different apico-basal Z levels of the same confocal stack (OHC1-2 field at the cochlear base). F-actin labels stereocilia and Afadin (AFDN) labels the apical junctions. In *Mpdz* mutants, GNAI enrichment is generally reduced, and often delocalized medially at the flat HC apex (arrows) and partially lost in the stereocilia (solid arrowheads). GNAI is also ectopically detected at medial apical junctions below the flat apex (hollow arrowheads). **D)** GNAI enrichment across the bare zone in P4 OHCs. The schematic depicts GNAI enrichment compared to the HC junction and the hair bundle in control and *Mpdz* mutant. The OHCs apical surface was divided in 3 regions of equal width (a, b, c) and GNAI signal was measured by region. N = number of animals, n = number of OHCs. Wilcoxon signed-rank test (*Mpdz^DEL/+^:* a vs b p= 0.1307 ^ns^; b vs c p= 0.3707 ^ns^; a vs c p= 0.9525 ^ns^; *Mpdz^DEL/DEL^:* a vs b and b vs c p< 0.0001 ****; a vs c p= 0.7880 ^ns^). **E)** GNAI enrichment at higher magnification in two representative P4 OHCs and IHCs (cochlear base; maximal projection). Note how in *Mpdz* mutant HCs, GNAI signal is reduced, particularly at the flat HC apex. In OHCs, GNAI is only retained in a subset of peripheral stereocilia (arrowheads) adjacent to where GNAI is retained at the peripheral region of the bare zone (arrows). Alternatively, GNAI is delocalized medially at the HC flat surface (orange arrows) and missing in the hair bundle entirely (orange arrowheads). **F)** Co-immunolabeling of GNAI (green) and acetylated tubulin (ACTUB, red) to reveal the kinocilium at P4 (cochlear base; maximal projection). A single OHC2 is outlined in each panel. As in control HCs, the kinocilium (arrowheads) is localized lateral to the hair bundle in *Mpdz* mutant OHCs, even when GNAI is delocalized medially (arrows). **G)** GNAI-GPSM2 co-labeling in P4 OHCs (cochlear base). Delocalized GNAI and GPSM2 (arrows) still co-localize in *Mpdz* mutants. Scale bars are 10μm (A), 5μm (C and E-G).

We next attempted to define more precisely how GNAI was redistributed in *Mpdz* mutant HCs. In P4 mutants OHCs at the base, GNAI was variably depleted at the bare zone and at stereocilia tips, and was instead ectopically found 1) on the longitudinal or medial sides of the hair bundle (Fig. 2C; arrows), and 2) at the medial HC junction with the neighboring support cell labeled with Afadin/AFDN (Fig. 2C; hollow arrowheads). Interestingly, GNAI was consistently missing at the center of the bare zone, whereas it was often retained at the bare zone periphery in *Mpdz* mutants. We quantified this trend by measuring GNAI enrichment in 3 equal sub-regions of the bare zone along the longitudinal axis (Fig. 2D). GNAI signals were equal in all sub-regions in control animals, as expected based on uniform GNAI enrichment across the bare zone. In contrast, *Mpdz* mutants showed a significant GNAI decrease in the central (b) compared to the peripheral sub-regions (a, c) (Fig. 2D). These results suggest that GNAI may be first lost centrally near the basal body, and invading the medial apical membrane by “seeping” through the sides of the hair bundle.

Interestingly, GNAI presence or absence at stereocilia tips coincided spatially with GNAI presence or absence at the adjacent portion of the bare zone in single OHCs (Fig. 2E). When GNAI was only maintained at the bare zone periphery (white arrow), GNAI was only maintained at the tips of peripheral stereocilia in the hair bundle (Fig. 2E, white arrowhead). Complete medial relocalization of GNAI outside the hair bundle (orange arrow) was paired with an almost complete loss of GNAI at tips. GNAI-GPSM2 enrichment at the bare zone is progressively reduced during HC maturation [8]. Low amounts of GNAI at the bare zone in P4 IHCs made delocalization to the flat surface and pairing with outcomes at tips challenging to assess in IHCs. We did however observe reduced GNAI bare zone enrichment in *Mpdz* mutant IHCs compared to controls, and a similar but less severe loss of GNAI at stereocilia tips compared to OHCs (Fig. 2E). Spatial pairing of GNAI between HC compartments in *Mpdz* mutants supports our working model where prior GNAI-GPSM2 enrichment at the bare zone instructs its selective trafficking to adjacent stereocilia in the first row [9, 10].

Because disrupted hair bundle morphology obscured hair bundle orientation in *Mpdz* mutants, we next immunolabeled the kinocilium at P4. As in control HCs, the kinocilium was systematically located on the lateral side of the hair bundle in *Mpdz* mutants (Fig. 2F, arrowheads). This clarifies that progressive delocalization of GNAI to the medial apex is not indicating a change in HC orientation. As expected, the GNAI binding partner GPSM2 followed GNAI delocalization in *Mpdz* mutants (Fig. 2G).

### The hair cell apical blueprint is globally disrupted in absence of MPDZ

INSC-GNAI-GPSM2 at the bare zone antagonizes aPKC at the medial apical surface, and the boundary between these two apical membrane compartments coincides with the lateral edge of the forming hair bundle [6, 8, 11]. Similar to GNAI and GPSM2, aPKC lost its spatial restriction in *Mpdz* mutants. At the cochlear base at P4, aPKC was restricted to the medial HC apex in control OHCs and IHCs, but ectopically found across the whole apical surface in *Mpdz* mutants (Fig. 3A). Measuring aPKC enrichment on each side of the hair bundle confirmed that the ratio between the lateral bare zone and the medial side was significantly higher in mutants compared to controls (Fig. 3B). This reflects an increase of aPKC intensity at the bare zone and a decrease at the medial side (see Supp. Data File). Of note, ectopic aPKC was previously reported upon *Gnai* and *Gpsm2* inactivation [6, 8], suggesting that aPKC invasion of the lateral apex in *Mpdz* mutants might be secondary to the downregulation of GNAI-GPSM2 there (Fig. 2). In contrast, aPKC was restricted medially as in control HCs at the cochlear apex at P4 (Fig. 3C), and along the whole cochlea at P0 (not shown).

**Figure 3.**
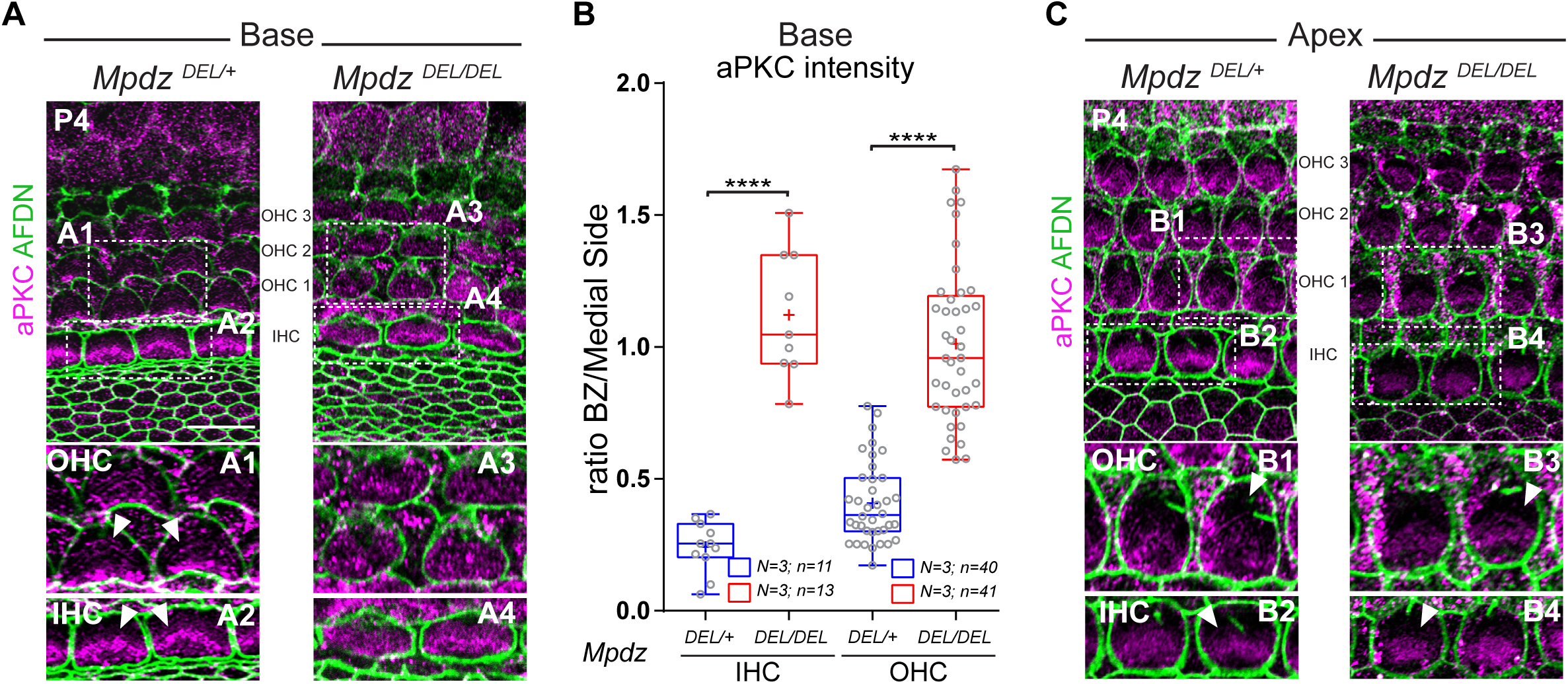
aPKC medial restriction at the apical membrane is lost in absence of MPDZ. aPKC labeling (magenta) at P4 at the cochlear base (A) and apex (C). Apical junctions are labeled with Afadin (AFDN; green). **A)** The lateral bare zone devoid of aPKC signal in controls (arrowheads) shows ectopic aPKC signal in *Mpdz* mutants in both OHCs and IHCs. **B)** aPKC enrichment at the HC apical surface at the cochlear base. The ratio between signal intensity at the bare zone (BZ) and at the medial side is plotted (see Supplementary Data File for values by compartments). N = number of animals, n = number of HCs (Mann-Whitney U test; OHC and IHC: p<0.0001 ****). **C)** At the cochlear apex, aPKC is normally excluded from the bare zone in *Mpdz* mutants. The boxed regions (A1-A4; B1-B4) are magnified in the bottom panels. Scale bars are 10μm.

Altogether, our results demonstrate that MPDZ is essential to maintain properly polarized and segregated GNAI-GPSM2 and aPKC domains in maturing HCs after P4. Delocalized GNAI-GPSM2 enrichment at the flat HC surface and its corresponding loss at stereocilia tips likely contribute to disrupted stereocilia alignment and generally dysmorphic hair bundles (Fig. 1-C).

### MPDZ defines a Crumbs complex at the apical membrane in hair cells and choroid plexus cells

MPDZ was previously reported to localize and function at tight junctions in epithelial cells [23, 35, 37]. Instead, our results revealed a role for MPDZ at the HC apical membrane, past the apical junction separating HCs from their support cell neighbors. MPDZ is a paralog of INADL/PATJ [37], a member of the CRB3-MPP5/PALS1-INADL Crumbs complex well known to regulate epithelial development and apico-basal polarization [26, 32, 33, 50]. We next compared the localization of MPDZ and other Crumbs complex members in neonate HCs using ZO1 as marker for apical junctions, and surface (XY) and projection views (XZ) of confocal stacks.

At P0 and P4, MPP5 was most highly enriched at the bare zone but also detected medially with little signal at the hair bundle (Fig. 4A), a distribution virtually identical to MPDZ (Fig. 1A). In addition, MPP5 was also detected at the HC apical junction (Fig. 4A, arrowhead). Projection views confirmed that MPP5, but not MPDZ, colocalized with ZO1 at OHC-support cell junctions (Fig. 4B, arrowheads). Occasional fixation artifacts separating the plasma membrane of HCs and neighboring support cells suggested that MPP5 was junctional in both cell types (Supp. Fig. 1A), as also expected from junctional MPP5 distribution at support cell-support cell contacts and outside the auditory epithelium (Fig. 4A). We tested multiple antibodies and protocols to immunolabel CRB3, the small integral member of the Crumbs complex. After repeated failures, we obtained decent results using an antibody against an intracellular CRB3 epitope and antigen retrieval with urea [51]. We observed CRB3 signal at the bare zone and at apical junctions, as for MPP5 (Fig. 4C). In contrast to other members of the Crumbs complex, INADL was not detected at the HC apical membrane or at the HC apical junction (Fig. 4D), and appeared instead limited to the apical junction of support cells (Supp. Fig. 1B). Together, these results suggest that a MPDZ-MPP5-CRB3 complex specifically occupies the HC apical membrane.

**Figure 4.**
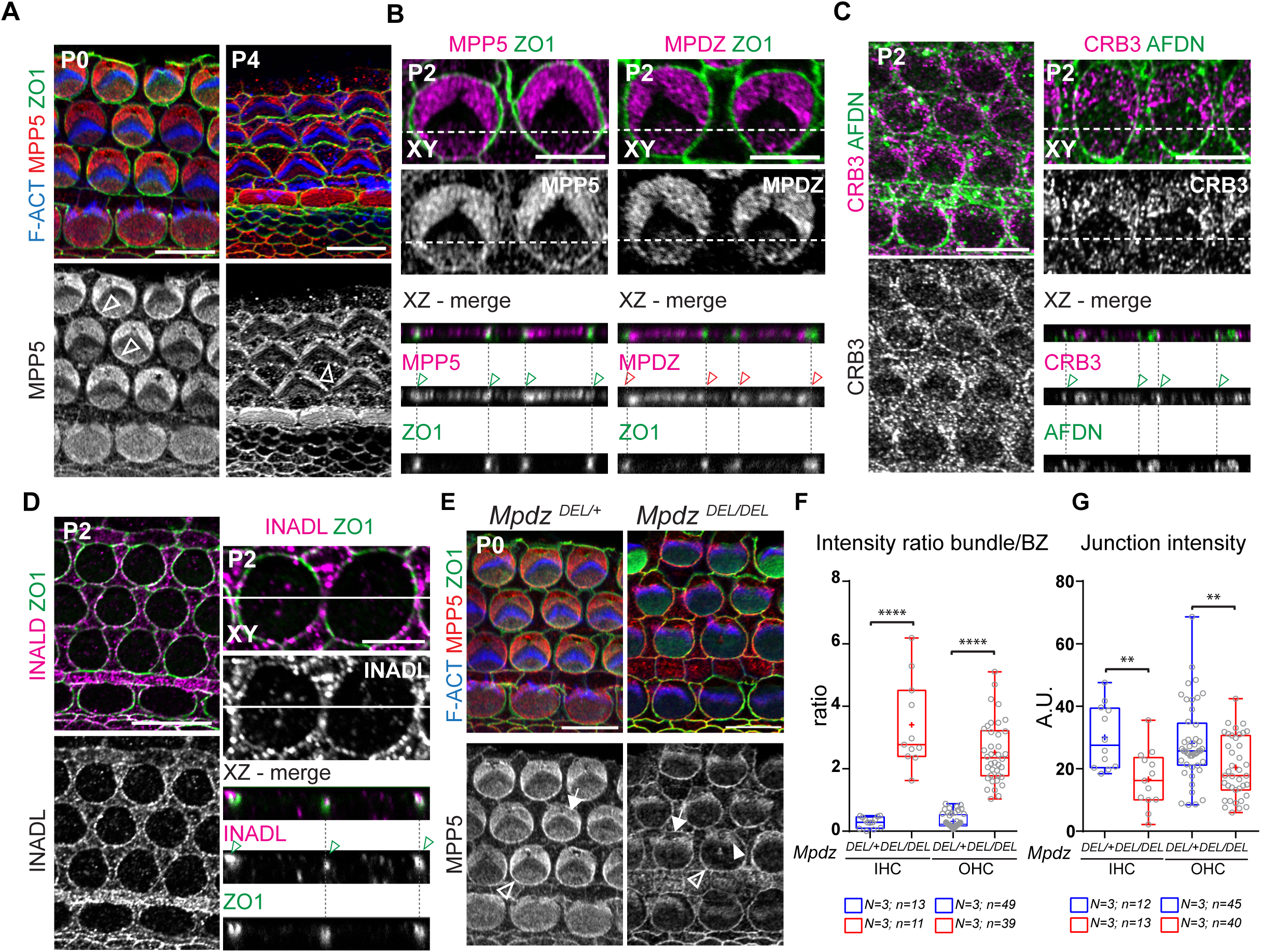
MPDZ coincides with MPP5/PALS1 and CRB3 and is required for their correct distribution at the hair cell apical membrane. **A-D)** Immunolabeling in flat (XY) and projection (XZ) views of the auditory epithelium between P0 and P4. **A)** MPP5 immunolabeling reveals a similar protein distribution to MPDZ (Fig. 1A). **B)** Projections also show MPP5 co-enrichment with ZO1 at the HC-support cell junctions (green arrowheads). MPP5 is enriched on both the HC and support cell membrane (see Supp. Fig. 1A). In contrast, MPDZ is not co-detected with ZO1 at apical junctions (red arrowheads). **C)** CRB3 distribution is similar to MPP5 (A-B), encompassing both the apical membrane and the apical junctions (green arrowheads). Afadin/AFDN was used to visualize apical junctions as ZO1 signal was compromised by the antigen retrieval procedure. **D)** INADL/PATJ is only detected at apical junctions with ZO1 (green arrowheads), and not at the apical surface. INADL appears to be enriched only on the support cell membrane (see Supp. Fig. 1B). **E)** MPP5 labeling in P0 *Mpdz* mutant. In absence of MPDZ, MPP5 is lost at the bare zone (arrow) and ectopically found in the hair bundle (solid arrowhead). MPP5 is retained at apical junctions (hollow arrowheads). **F-G)** Apical MPP5 enrichment in P0 HCs at the cochlear base. The ratio between the MPP5 signals at the hair bundle and bare zone (F) and the MPP5 signal at the medial apical junction (G) are plotted. N = number of animals, n = number of HCs (Mann-Whitney U test; (F) IHC and OHC: p<0.0001 ****; (G) IHC: p=0.00086 **, OHC p=0.0079 **). Cochlear base (A, C left, D left, E), cochlear mid (B, C right, D right). Scale bars are 10μm.

Unlike MPDZ, MPP5-CRB3 distribution also included apical junctions in HCs, and loss of MPDZ was proposed to disrupt barrier integrity in ependymal and choroid plexus cells in part by compromising tight junctions [41, 42]. We thus immunolabeled the choroid plexus (4^th^ ventricle) in P13-P15 wild-type mice. Interestingly, we found that, as in the auditory epithelium, MPDZ was absent at ZO1-positive cell junctions but found at the apical membrane with higher enrichment near the junctions (Supp. Fig. 2A, E). In contrast, MPP5 and CRB3 were clearly detected at apical junctions in addition to the apical membrane, as in HCs (Supp. Fig. 2B-C). INADL was not detected at all (Supp. Fig. D). We conclude that in auditory and choroid plexus epithelial cells at least, MPDZ-MPP5-CRB3 represents a Crumbs complex specifically addressed to the apical membrane. MPP5-CRB3 also occupies apical junctions, potentially with another partner.

### MPDZ anchors MPP5 and CRB3 at the hair cell apex

MPDZ is expected to function as an anchor for partner proteins carrying a PDZ-binding domain, and MPDZ and MPP5 are known to interact directly [37, 52]. We wondered whether the loss of MPDZ would disrupt Crumbs complex partners in addition to GNAI-GPSM2 (Fig. 2) and aPKC (Fig. 3). At P0, MPP5 was missing at the bare zone in *Mpdz* mutants, and apparently ectopically found in the hair bundle (Fig. 4E; solid arrowhead). Measuring MPP5 intensity ratio between the bare zone and the hair bundle confirmed this striking redistribution in *Mpdz* mutants (Fig. 4F). In contrast, MPP5 was still enriched at HC junctions in *Mpdz* mutants (Fig. 4E; hollow arrowheads), in line with our inability to detect MPDZ at junctions (Fig. 4B). Measuring MPP5 junctional signal revealed a modest reduction in *Mpdz* mutants (Fig. 4G), but this could reflect the difficulty to quantify junctional signals without including signals at the adjacent apical membrane. Similarly, we observed that CRB3 signal was lost at the bare zone and ectopically found in the hair bundle in P0 *Mpdz* mutants (Supp. Fig. 1C). As expected from its localization at support cell junctions, INADL was unchanged in *Mpdz* mutants (Supp. Fig. 1D).

As we observed earlier disruption of MPP5 and CRB3 (P0) than GNAI-GPSM2 and aPKC (P4) in *Mpdz* mutants, we conclude that MPDZ is directly required to anchor its Crumbs complex partners MPP5-CRB3 at the HC apical membrane. In turn, loss of the Crumbs complex at the flat apical membrane disrupts the polarization and segregation of GNAI-GPSM2 and aPKC.

### GNAI-GPSM2 and MPDZ-MPP5 influence each other at the hair cell apical membrane

MPDZ is required to maintain the lateral polarization of GNAI-GPSM2 at the apical membrane from P4 (Fig. 2), and MPDZ-MPP5-CRB3 is more highly enriched at the bare zone compared to the medial HC surface (Fig. 1A, Fig. 4A, C). We thus reasoned that, reciprocally, GNAI-GPSM2 might influence the distribution of MPDZ and MPP5 by increasing their accumulation at the bare zone. To test this prediction, we analyzed a constitutive *Gpsm2* mutant (*Gpsm2^DEL^*) [8] and we globally downregulated GNAI activity in HCs by expressing the catalytic subunit of Pertussis toxin (PTXa) with *Atoh1-Cre* [9]. At P4, *Gpsm2* mutant HCs lost preferential MPDZ enrichment at the bare zone (Fig. 5A). Measuring the ratio of signal intensity on the medial versus lateral side of the hair bundle confirmed a more uniform MPDZ distribution across the apical surface upon loss of GPSM2 (Fig. 5B). Surprisingly however, these quantifications also revealed that MPDZ signal intensity actually increased both on the medial and lateral sides in *Mpdz* mutants, only more drastically on the medial side (Fig. 5C).

**Figure 5.**
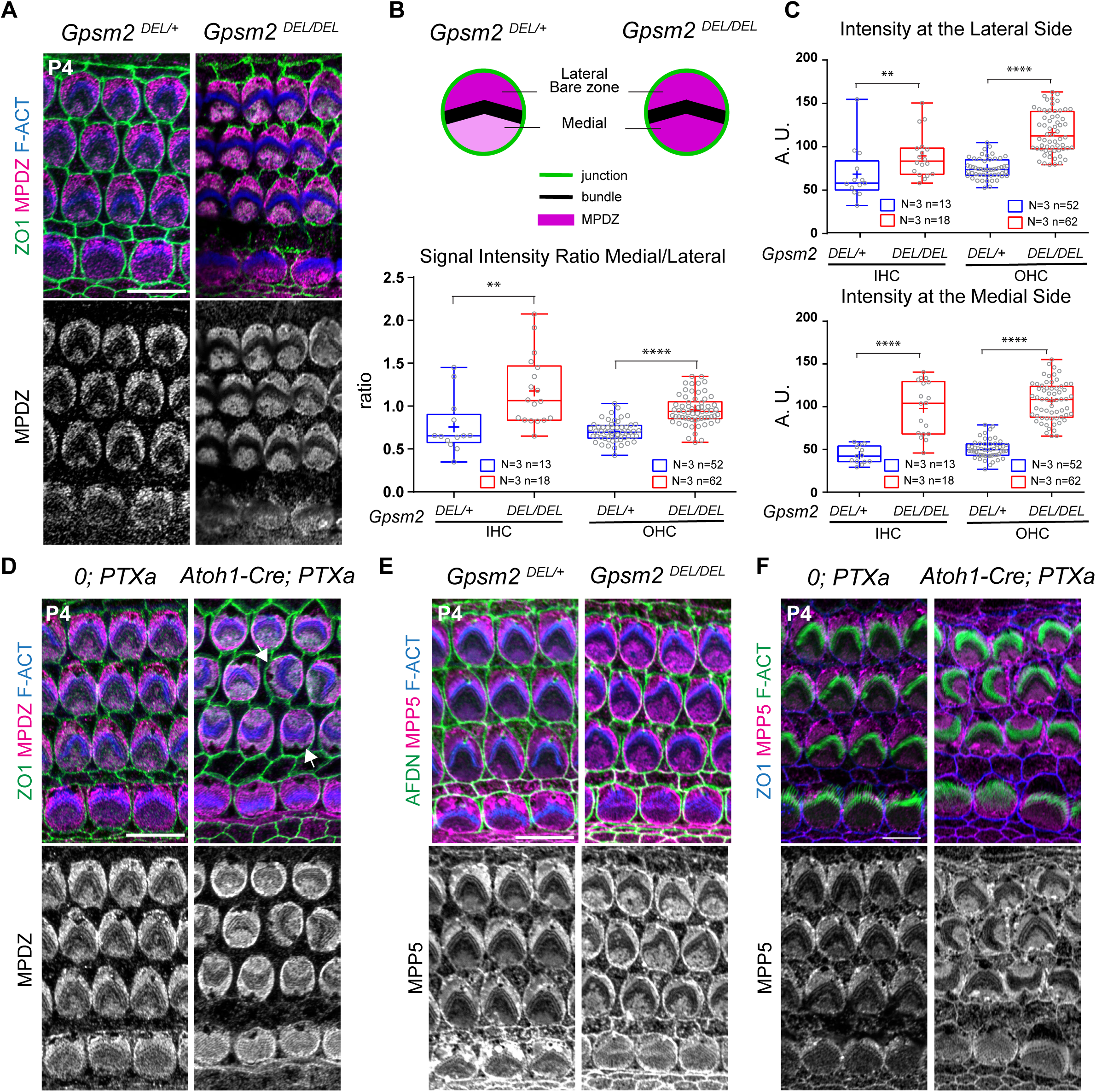
GNAI-GPSM2 promotes planar polarization of MPDZ and MPP5 at the apical membrane. **A)** MPDZ labeling (magenta) in *Gpsm^DEL/DEL^* HCs at P4 (cochlear apex). MPDZ lacks higher enrichment at the lateral bare zone in absence of GPSM2. **B-C)** MPDZ enrichment in *Gpsm2* mutant HCs at P4 (cochlear apex). The ratio between the medial and lateral (bare zone) MPDZ signals (B), and the individual signals at the bare zone (C, top) and medial apex (C, bottom) are plotted. Note how MPDZ signals increase in both compartments in *Gpsm2* mutants, but more strongly at the medial apex. N = number of animals, n = number of HCs (Mann-Whitney U test; (B) IHC: p=0.0014 ***, OHC: p<0.0001****; (C) Lateral IHC: p=0.0064 **, OHC: p<0.0001 ****; Medial IHC and OHC: p<0.0001 ****). **D)** MPDZ labeling (magenta) in Pertussis toxin (PTXa)-expressing HCs at P4 (cochlear apex). MPDZ lacks higher enrichment at the lateral bare zone when GNAI is inactivated by PTXa. GNAI inactivation provokes graded OHCs misorientation so that the bare zone region (arrows) is not consistently lateral in PTXa OHCs. **E-F)** MPP5 labeling (magenta) in *Gpsm^DEL/DEL^* (E) and PTXa (F) HCs at P4 (cochlear apex). Like MPDZ (A-D), MPP5 is more uniformly enriched across the apical membrane when GPSM2 or GNAI is inactivated. In (E), Afadin (AFDN) is used instead of ZO1 to label the apical junctions. Scale bars are 10μm.

Relatedly, we observed more uniform MPDZ distribution across the HC apex in PTXa-expressing HCs at P4 (Fig. 5D). Of note, PTXa provokes severe defects in OHC orientation [8, 9], and consequently the bare zone was no longer systematically on the lateral side of the auditory epithelium. We next immunolabeled MPP5 in *Gpsm2* and PTXa mutants since MPP5 binds [37, 52] and co-localized with MPDZ at the HC surface. As with MPDZ, preferential MPP5 enrichment at the bare zone was abolished in *Gpsm2* (Fig. 4E) and PTXa (Fig. 5F) mutant HCs at P4. Altogether, our results suggest that reciprocal interactions between MPDZ-MPP5 and GNAI-GPSM2 ensure that proper amounts of each protein complex are maintained at the bare zone. Increased MPDZ enrichment on both sides of the hair bundle in absence of GPSM2 may indicate that GNAI-GPSM2 somehow limits the total amount of MPDZ allowed to occupy the apical membrane.

### Progressive loss of the apical blueprint coincides with severe hair bundle defects and hearing loss in mature hair cells

Early loss of GNAI and GPSM2 in postmitotic HCs leads to hair bundle disorganization, stereocilia stunting and profound hearing loss in young adults [5, 7, 9, 10]. We asked how loss of MPDZ that provokes a later disruption of GNAI-GPSM2 (Fig. 2) impacts the auditory organ near maturity. At P21, we did not observe OHC or IHC death along the cochlea in *Mpdz* mutants based on MYO6 immunolabeling (Fig. 6A). This verifies that delocalized GNAI-GPSM2 and aPKC starting at P4 (Fig. 2–3) is not an indirect consequence of HC degeneration, but an active process.

**Figure 6.**
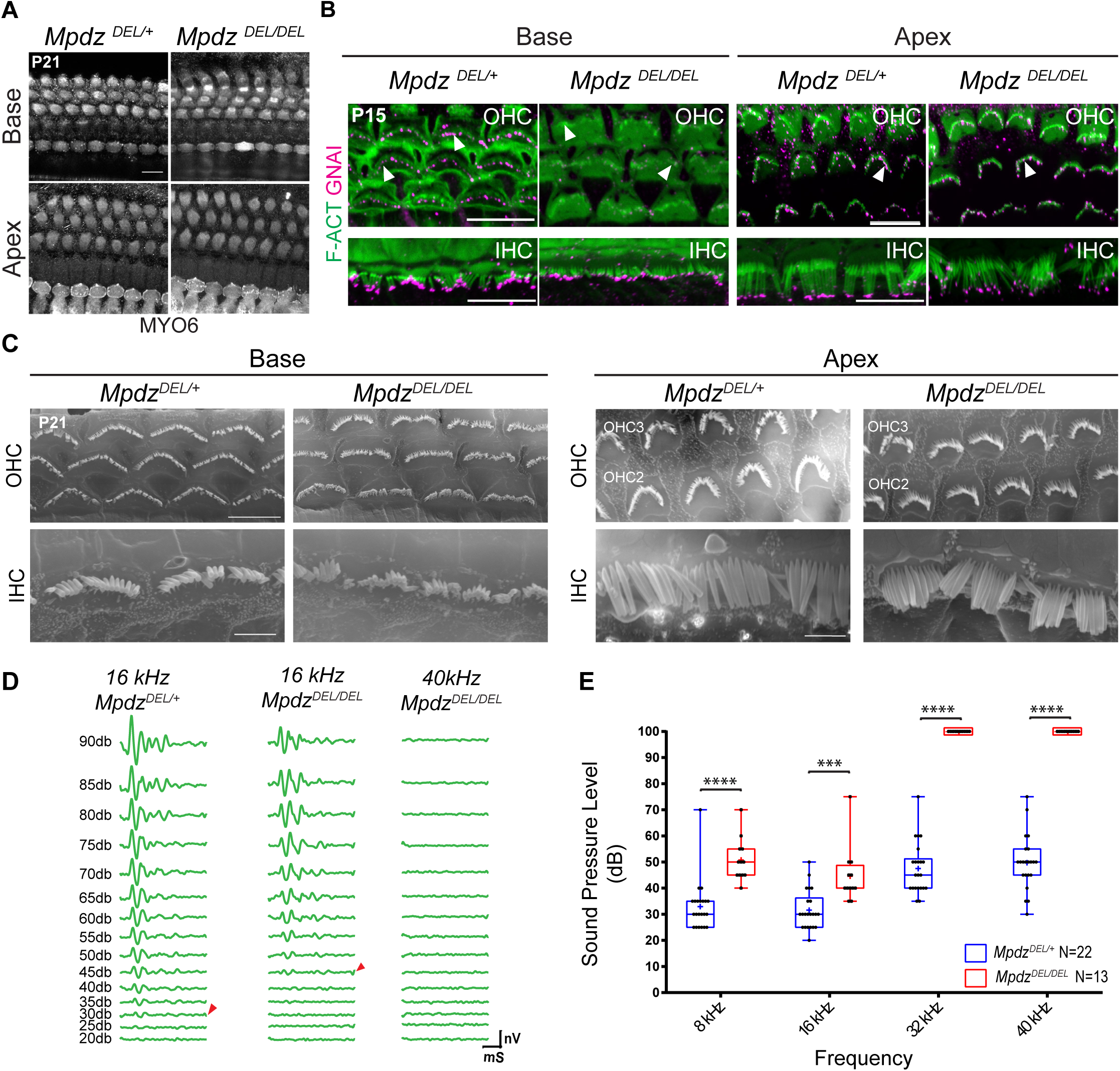
Loss of MPDZ leads to tonotopically matching hair bundle and hearing defects in young adults. **A)** MYO6 labeling of the auditory epithelium at P21 (maximal projection). In spite of severe apical patterning defects in neonates, no HC death is observed in young adults. **B)** GNAI labeling (magenta) at P15 (maximal projection). GNAI signal at stereocilia tips (arrowheads) in control OHCs is lost in *Mpdz* mutant at the cochlear base, but retained at the cochlear apex. In IHCs, GNAI signal is mostly retained at tips even at the cochlear base. **C)** Scanning electron microscopy images of representative OHCs and IHCs at P21 at the cochlear base and apex. Defects include flat and misshapen OHC bundles, and missing or irregular stereocilia placement in the first row in IHCs at the cochlear base. Hair bundle morphology in OHCs and IHCs is relatively normal at the cochlear apex. **D)** Auditory brainstem response (ABR) waveforms at variable sound pressure levels for one control littermate (16kHz) and one *Mpdz* mutant (16 and 40 kHz). The threshold of ABR response is indicated (arrowheads). **E)** ABR thresholds by genotype (8, 16, 32, 40 kHz). Animal were tested between P21 and P25. N = number of animals (two-way ANOVA test with a multiple comparison test; 8kHz: p<0.0001 ****; 16kHz: p<0.001 ***; 32kHz: p<0.0001 ****; 40kHz: p<0.0001 ****). A plotted value of 100dB indicates that animals did not respond at 90bD. Scale bars are 10μm (A-B, C for OHCs), 5μm (C for IHCs).

GNAI delocalization in *Mpdz* mutants progresses from the cochlear base to the apex along with HCs differentiation (Fig. 2A). We immunolabeled GNAI at P15 to determine how this defect progresses beyond P7 (Fig. 2A). At the cochlear base, GNAI signal was visible at stereocilia tips in controls, but largely absent in mutant OHCs (Fig. 6B). In contrast, GNAI could be observed at stereocilia tips in both control and mutant OHCs at the cochlear apex (Fig. 6B). We conclude that variable loss of GNAI at stereocilia tips at P4 (Fig. 2E) evolves into a complete loss with time, and that the GNAI delocalization wave (Fig. 2A) never reaches the apex of the cochlea in *Mpdz* mutants. In P15 IHCs, GNAI signal at stereocilia tips was still largely visible both at the cochlear base and apex in *Mpdz* mutants (Fig. 6B). This suggests that the milder GNAI disruption observed at P4 in IHCs (Fig. 2E) does not evolve much by P15, or perhaps that GNAI trafficking to IHC tips is only delayed in *Mpdz* mutants.

To better resolve the hair bundle structure, we performed scanning electron microscopy (SEM) at P21. At the cochlear base, mutant OHCs retained a flattened hair bundle and apical surface circumference (Fig. 6C), as seen in neonates (Fig. 1B). Mutant IHCs were less affected but showed irregular stereocilia placement (Fig. 6C). At the cochlear apex by contrast, OHC and IHC morphology was relatively normal, in line with normal localization of GNAI at the bare zone (P7, Fig. 2A) and stereocilia tips (P15, Fig. 6B).

To pair morphological alterations with organ function, we recorded Auditory Brainstem Response (ABR) in 4 week-old *Mpdz* mutants and control littermates. We presented anesthetized animals with pure tone stimuli of variable sound pressure intensities (dB) to record ABR waveforms with a subcutaneous probe (Fig. 6D). Plotting ABR thresholds (Fig. 6D, arrowheads) by frequency and genotype revealed severe global hearing defect in *Mpdz* mutants (Fig. 6E). At high frequencies (32 and 40 kHz), *Mpdz* mutants showed complete hearing loss, with virtually no response to even the loudest stimuli (90dB). At lower frequencies (8 and 16 kHz), *Mpdz* mutants did respond, but with a significantly higher threshold compared to control littermates. More severe hearing loss at higher frequencies was expected based on the tonotopic organization of HC frequency sensitivity along the cochlear duct. HCs at the cochlear apex, less affected in *Mpdz* mutants, detect lower frequencies.

## DISCUSSION

We show that the polarity protein MPDZ is required to anchor a version of the Crumbs complex (MPDZ-MPP5-CRB3) at the HC apical membrane. Loss of MPDZ (and concomitant defects in MPP5-CRB3 apical enrichment) disrupts the maintenance of GNAI-GPSM2 and aPKC in their polarized and segregated apical domains. Blurring of this molecular blueprint must contribute to stereocilia misalignment and dysmorphic hair bundles in *Mpdz* mutants, as similar defects were reported in absence of GNAI or GPSM2. In turn, defective hair bundle morphogenesis is probably a leading cause for hearing loss in *Mpdz* mutants.

In absence of MPDZ, hair bundle defects are visible at birth already, whereas defects in the GNAI-GPSM2 vs aPKC apical blueprint and OHC circumference are only noted from P4. While delocalization of GNAI-GPSM2 is likely an important contributing factor, loss of MPDZ must interfere with other proteins required for early hair bundle development. Interestingly, *Mpdz* mutant HCs show a striking relocalization of both MPP5 and CRB3 from the flat HC apex to the hair bundle at P0 (Fig. 4E; Supp. Fig. 1C). CRB3 was shown to interact with actin-binding proteins including EPS8 [53] and Ezrin [33]. It is thus possible that ectopic MPP5-CRB3 presence in growing stereocilia interferes with F-actin assembly or with the association between F-actin and the plasma membrane, for example.

Of note, progressive delocalization of GNAI-GPSM2 and aPKC at the HC apical membrane was previously reported in mutant HCs lacking the SMARCA4/BRG1 subunit of the SWI/SNF chromatin remodeling complex [54]. Loss of SMARCA4 protein between P1-P4 caused HCs to start dying from P8 however, questioning whether loss of protein polarization at the HC apical membrane might possibly be a side effect of cellular decline. In contrast to SMARCA4, loss of MPDZ does not lead to gross epithelial disorganization, and all HCs survive at least until P21. Maintaining the molecular blueprint that regulates cytoskeleton distribution at the HC surface is thus an active process. This conclusion is supported by another recent study where disrupting proteostasis in postnatal HCs also affected the polarized distribution of blueprint proteins in absence of HC death [55].

*Mpdz* is one of few genes associated with congenital hydrocephalus, and two studies inactivating *Mpdz* in mouse reported severe hydrocephalus from P3 that proved lethal before, or around 3 weeks of life [41, 42]. The mouse models analyzed were a gene-trap allele in intron 11-12 [41, 42, 46] and a conditional deletion of exons 4-5 [41], both expected to produce a loss-of-function. Intriguingly, we do not observe hydrocephalus in *Mpdz^DEL/DEL^* mutants produced by the KOMP consortium (C57BL/6N-*Mpdz^em1(IMPC)J^*/Mmucd), where an indel in exon 6 is expected to truncate the protein after the first of 13 PDZ domains. We observed a sub-mendelian frequency of homozygotes before weaning age, but we could obtain and breed *Mpdz^DEL/DEL^* adults of both sexes (43% of progeny from *Mpdz^DEL/DEL^* x *Mpdz^DEL/+^* was homozygote; see Supp. Table 1 for a comparison between *Mpdz* alleles). The cause of this discrepancy is unclear, but may involve differences in genetic background between the C57BL6/J and C57BL6/N strains. Study of another gene linked to congenital hydrocephalus, *L1cam*, revealed a potent modifier locus on chromosome 5 [56]. Importantly, we use a validated MPDZ antibody [34] that showed a loss of signal in the choroid plexus of gene-trap *Mpdz* mutants [42]. A similar absence of signal in *Mpdz^DEL/DEL^* HCs (Figure 1A) thus validates the KOMP allele as a bona fide loss-of-function. This conclusion is reinforced by severe morphological and auditory defects in *Mpdz^DEL/DEL^* mutants.

Interestingly, MPDZ is limited to the apical membrane in HCs and choroid plexus epithelial cells, and could not be detected along with MPP5-CRB3 at apical junctions. Loss of MPDZ does however affect the structure of apical junctions in choroid plexus epithelial cells [42], and the circumference of OHCs in the auditory epithelium (Fig. 1D-E). The latter might be an indirect consequence where hair bundle defects affect HC junction remodeling [48], but it is not unexpected that apical membrane proteins in the vicinity of tight junctions would participate in the establishment and maintenance of junctional integrity. In fact, a recent study identified the Crumbs complex (INADL-MPP5-CRB3) as defining a transition zone between tight junctions and the free apical membrane, which the authors coined as the ‘vertebrate marginal zone’ [57].

In other contexts by contrast, MPDZ was reported as enriched at apical junctions [23, 35, 37–39]. For example, the G protein regulator DAPLE and MPDZ co-localize and cooperate at junctions during apical cell constriction in cultured cell monolayers and in neuroepithelial cells [38, 39]. Moreover, *DAPLE* is with *MPDZ* one of a few genes linked to congenital hydrocephalus in human and mice [58–60]. In auditory HCs however, DAPLE is enriched laterally at the apical HC junction [18], a distinct distribution from MPDZ at the HC apical membrane. In addition, apical HC defects in *Daple* mutants are more severe and distinct from the ones we report here in *Mpdz* mutants [18]. We thus conclude that the epithelial cell context must dictate what binding partners and subcellular compartment MPDZ adopts.

In summary, our results show that MPDZ specifically patterns the HC apical membrane, in part by maintaining the proper planar segregation of blueprint proteins at neonate stages. Loss of MPDZ leads to permanent hair bundle disorganization that is likely the etiology of severe hearing loss in young adults.

## ACKNOWLEDGMENTS

We are particularly grateful to Dr. Elior Peles (Weizmann Institute of Science) for sharing the MPDZ antibody, and to Dr. Xaralabos Varelas (Boston University) for sharing the CRB3 antibody. The *Mpdz* mouse model was produced by the KOMP program led by Steve Murray at The Jackson Laboratory, and was shared by the group of Patsy Nishina. A.J. was supported by a 2-year postdoctoral fellowship from Fondation pour l’Audition, Paris, France (FPA RD2018-3). This work was supported by the National Institute on Deafness and Other Communication Disorders with R01 DC015242 and DC018394 to B.T.

## AUTHOR CONTRIBUTION

B.T conceived the study. A.J and B.T. designed, and A.J performed and analyzed all experiments. B.T. and A.J. wrote the manuscript, B.T. secured funding.

## COMPETING INTERESTS

The authors declare no competing interests.

## DATA AVAILABILITY STATEMENT

All data generated or analyzed in this study are included in this published article (and its supplementary information file).

## METHODS

### Mice

The *Mpdz^DEL^* strain (C57BL/6N-*Mpdz^em1(IMPC)J^*/Mmucd, MGI: 5571597) was produced by the KnockOut Mouse Project Consortium (KOMP) at The Jackson Laboratory. *Gpsm2^DEL^ (Gpsm2^tm1a(EUCOMM)Wtsi^*; MGI:4441912 [8]) and *R26-loxP-stop-loxP-mycPTXa* (*Gt (ROSA)26Sor^em1(ptxA)Btar^*; MGI:6163665 [9]) were described previously. Wild-type samples analyzed were in a mixed genetic background (C57BL/6J x FVB/NJ). Mouse samples analyzed ranged in age from E17.5 to 4-week old as indicated in each figure, and included both males and females. Heterozygote *Mpdz*^*DEL*/+^ animals did not exhibit morphological, molecular or hearing defects, and were thus systematically used as littermate controls for *Mpdz^DEL/DEL^* mutants to reduce the number of animals produced for the study. Animals were maintained under standard housing and all animal work was reviewed for compliance and approved by the Animal Care and Use Committee of The Jackson Laboratory.

### Immunofluorescence and antibodies

For embryonic and neonate stages, temporal bones were isolated and either immediately micro-dissected to expose the sensory epithelium before fixing in paraformaldehyde (PFA 4%) for 1h at 4°C, or incubated in trichloroacetic acid (TCA 10%) for 10min on ice before dissection, depending on the antibodies used (indicated below). After rinsing with PBS, the tectorial membrane was removed, and samples were blocked and permeabilized in PBS with 0.5% Triton-X100 and bovine serum albumin (1%) for at least 1h at room temperature before application of the primary antibodies. For CRB3 labeling, we used the following antigen retrieval procedure: the postnatal inner ear was fixed before micro-dissection for 1h at room temperature in PFA. The auditory epithelium was then exposed, blocked and permeabilized as above for 20 minutes, and the incubated in urea (0.8M) for 1h at 50°C, then for 20 minutes at 65°C.

For postnatal stages after P7, temporal bones were isolated and the cochlea punctured at the apex to facilitate access of the fixative before immersion fixation in paraformaldehyde (PFA 4%) for 1h at 4°C or room temperature. After rinsing with PBS, the temporal bone was incubated overnight at room temperature in 0.11M EDTA for decalcification before dissection to expose the auditory epithelium. Samples were then blocked and permeabilized following the same protocol as for earlier stages.

The hindbrain (4^th^ ventricle) choroid plexus was dissected at P13 or P15, and fixed in paraformaldehyde (PFA 4%) for 1h at 4°C. After rinsing with PBS, samples were blocked and permeabilized in PBS with 0.5% Triton-X100 and bovine serum albumin (1%) for at least 1h at room temperature before application of the primary antibodies.

Primary and secondary antibodies were each incubated overnight at 4°C in PBS. Fluorescent-conjugated phalloidin was added with the secondary antibodies. After each antibody incubation, samples were washed 3 times with PBS + 0.05% Triton-X100 before a final post-fixation in PFA 4% for 1h at room temperature. Samples were then mounted flat on a microscopy slide (Denville M1021), either directly under a 18×18mm #1.5 coverglass (VWR 48366-045) (cochlea), or using office tape as spacer (choroid plexus). Mowiol was used as mounting medium (Calbiochem/MilliporeSigma 4759041). Mowiol (10% w/v) was prepared in 25%(w/v) glycerol and 0.1M Tris-Cl pH8.5.

Primary antibodies used were:

rabbit anti-MPDZ [34] (gift from Elior Peles, Weizmann Institute of Science; PFA)
rabbit anti-CRB3 [51] (gift from Bob Varelas, Boston University; PFA)
rabbit anti-TRIOBP/TARA (Proteintech; 16124-1-AP; PFA)
rabbit anti-GNAI1/2/3 (Santa Cruz; sc-262; PFA; results in our laboratory indicate that this reagent is not specific for a GNAI paralog and that at least GNAI2 and GNAI3 are co-localized apically in hair cells. We thus use “GNAI” to refer to the signal obtained)
rabbit anti-GPSM2 (Sigma; A41537; PFA)
mouse anti-aPKC/PRKCZ (Santa Cruz; sc-216; PFA)
sheep anti-AFDN/AF-6 (R&D; AF7829; PFA)
rat anti-ZO1 (Developmental Studies Hybridoma Bank; R26.4C; TCA or PFA)
rabbit anti-MPP5 (Santa Cruz; sc-33831; PFA)
rabbit anti-INADL (LifeSpan Biosciences; LS-C410011-50; TCA)
rabbit anti-MYO6 (Proteus; 25-6791; PFA)
mouse anti-acetylated Tubulin (Santa Cruz; 23950; PFA)
Secondary antibodies were raised either in goat or donkey, and coupled to Alexa Fluor (AF) 488, 555 or 647 (ThermoFisher Scientific). We used fluorescent conjugated phalloidins to reveal F-Actin (ThermoFisher Scientific: AF488, A12379, Biotium: CF405, 89138-126).

### Image acquisition and quantification

Images were captured using a LSM800 line scanning confocal microscope using the Zen2.3 software, the Airyscan detector in regular confocal mode, and a 63x 1.4 NA oil objective lens (Carl Zeiss AG). When not stated otherwise in the figure legend, all images show a single optical plane. Images were processed using Adobe Photoshop (CC 2020), and the same image treatment was applied across genotypes or conditions in the same experiment. In all experiments, quantifications include three animals of each genotype. All experimental values plotted in the study, as well as animal cohort size and hair cell numbers are detailed in a single Supplementary Data File where each tabulation reports on a single experiment. Results in all 3 OHC rows were pooled because we did not observe a row-specific trend.

To measure hair cell apical surface area (Figure 1E), z-series stacks were acquired for each sample either at the base or the apex, and a single Z slice was selected based on junctional ZO1 signal. The ZO1-positive outline of the hair cell was traced and the surface area measured using Fiji/ImageJ. To measure immunolabeling signal intensity, microscopy fields were selected where hair cells were mounted as flat as possible so as to avoid artifacts. Z-series stacks were acquired at the cochlear base using the same laser intensity and gain for all samples (mutant and control littermates), and a single Z slice/optical plane was chosen where the signal was the strongest. ZO1 or Afadin/AFDN were used to label the apical hair cell junction, and the base of the stereocilia bundle (phalloidin) was used to distinguish the bare zone (lateral) and the medial sides of the apical membrane. Fiji/ImageJ was used to measure the Mean gray value in a region of interest (ROI) as defined below. For each image, background signal was sampled and averaged, then subtracted to all hair cell quantifications on the same image.

To measure GNAI signal intensity in *Mpdz* mutants (Figure 2D), the lateral (bare zone) and medial compartments were divided in 3 ROIs of equal width (a, b, c; see Figure 2D), and the Mean gray value of each region was measured. To obtain the Mean gray value of the entire bare zone and medial compartments (Figure 2B), values in the 3 ROIs were averaged. To measure aPKC (Figure 3B) and MPP5 (Figure 4F) signals in *Mpdz* mutants, a fixed ROI was positioned so as to cover about 50% of the apical compartments considered (bare zone, medial apex, hair bundle). To measure MPP5 intensity at junctions (Figure 4G), a rectangular ROI of 0.85 μm^2^ (1.67 × 0.51μm) was positioned at the medial hair cell junction. To measure MPDZ intensity in *Gpsm2* mutants (Figure 5B-C), a circular ROI of 1.054 μm^2^ was positioned at the lateral (bare zone) and medial apical compartments.

### Scanning Electron Microscopy (SEM)

Temporal bones were isolated from the skull, a hole was punctured at the cochlear apex, and samples were immersion-fixed for at least one overnight at 4°C in 2.5% glutaraldehyde + 4% paraformaldehyde (Electron Microscopy Science) in a 1mM MgCl2, 0.1M Sodium Cacodylate, 20mM CaCl_2_ buffer. After rinses in PBS, samples were decalcified overnight with 0.11M EDTA and the auditory epithelium was dissected in 3 pieces (cochlear base, mid, apex). Dissected samples were progressively dehydrated in an ethanol series (30-50-70-80-90-100%, for at least 20 minutes each). Chemical sample drying was done with hexamethyldisilazane (HMDS; Electron Microscopy Science 50-243-18). Dry samples were then mounted on aluminum stubs using double-sided carbon tape, and sputter-coated with gold-palladium before imaging on a Hitachi 3000N VP electron microscope. Images were taken at 20kV and with a 5k magnification.

### Auditory Brainstem Response (ABR) tests

Mice were anesthetized by intraperitoneal injection of a mix of ketamine and xylazine (10 mg and 0.1 mg per 10g of body weight, respectively) and body temperature was maintained at 37°C using a heating pad (FHC). All tests were conducted in a sound-attenuating chamber. ABR testing was done using the RZ6 Multi-I/O Processor System coupled to the RA4PA 4-channel Medusa Amplifier (Tucker-Davis Technology). Tone bursts at 8, 16, 32 and 40 kHz were generated with the TDT system, and sub-dermal needles were used as electrodes, with the active electrode inserted at the cranial vertex, the reference electrode under the left ear, and the ground electrode at the right thigh. Auditory thresholds were obtained for each stimulus by reducing the Sound Pressure Level (SPL) by 5 dB steps between 90 and 20 dB to determine at which SPL an ABR could be recognized. *Mpdz^DEL/DEL^* mutants and *Mpdz^DEL/+^* control littermates were tested between age P21 and P24. Both sexes were tested (see detailed Supplementary Data File), and because sex had no influence on thresholds, sexes were pooled in Figure 6E.

### Statistical Analysis

Data were plotted in Prism 8(GraphPad). All values were plotted individually, and the distribution was framed with 25-75% whisker boxes where exterior lines show the minimum and maximum, the middle line representing the median, and + is used for the mean. A potential difference in data distribution between genotypes was tested for significance using the Mann-Whitney U test (non-parametric unpaired t-test), except in the following cases: i) GNAI signal intensity by sub-region at the bare zone (Figure 2D), where we used a Wilcoxon signed-rank paired test to compare the 3 different sub-regions (a, b, c). ii) ABR thresholds (Figure 6D), where we used the 2-way ANOVA test coupled with a Sidak’s multiple comparison. Exact p-values are indicated in the figure legends, and p-values are summarized on graphical plots with the following “star”: system: p<0.0001 = ****, 0.0001<p<0.001 = ***, 0.001<p<0.01 = **, 0.01<p<0.05 = *, p>0.05 = not significant (ns). When not quantified, all immunolabeling experiments included at least 3 mutant samples in two litters, and a similar number of littermate control animals. In that case, figure panels show a representative outcome observed in all mutant samples.

## Supplementary Figures

**Supplementary figure 1.**
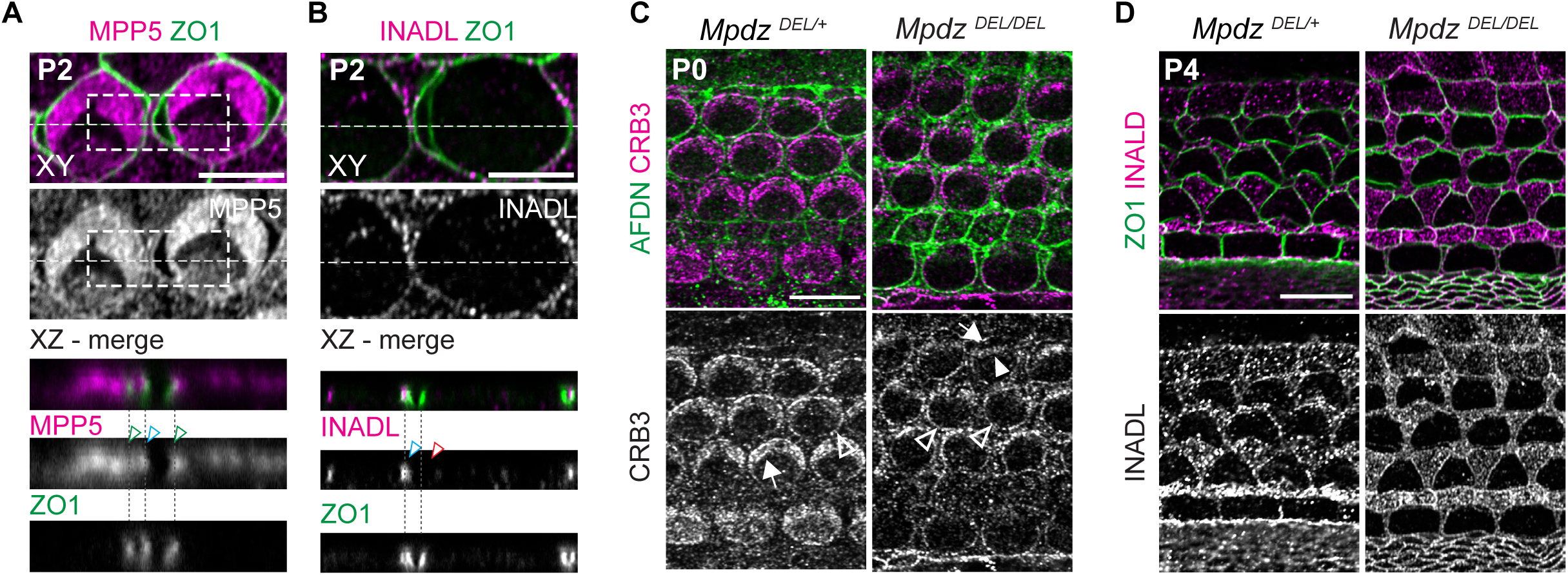
MPP5 and INADL are enriched at the apical junction in support cells. MPDZ is required for the correct distribution of CRB3 at the hair cell apical membrane. **A-B)** Immunolabeling in flat (XY) and projection (XZ) views of two OHC3 at the cochlear mid. We took advantage of TCA fixation artifacts where the apical membrane of the HC and neighboring support cell became separated. MPP5 (A) is enriched at both HCs (green arrowheads) and support cells (blue arrowheads) apical junctions. In contrast, INADL (B) is only enriched at the support cell apical junction (blue arrowhead) and absent at the HC junction (red arrowhead). **C)** CRB3 labeling in P0 *Mpdz* mutant (cochlear base). In absence of MPDZ, CRB3 is lost at the bare zone (arrow) and ectopically found in the hair bundle (solid arrowhead). CRB3 is retained at apical junctions (hollow arrowheads). Note how CRB3 behaves similarly to MPP5 (see Figure 4E-F). **D)** INADL labeling in P4 *Mpdz* mutant (cochlear base). INADL enrichment remains unchanged in absence of MPDZ, as expected from the absence of INADL signal in HCs (B; see also Figure 4D). Scale bars are 10μm.

**Supplementary figure 2.**
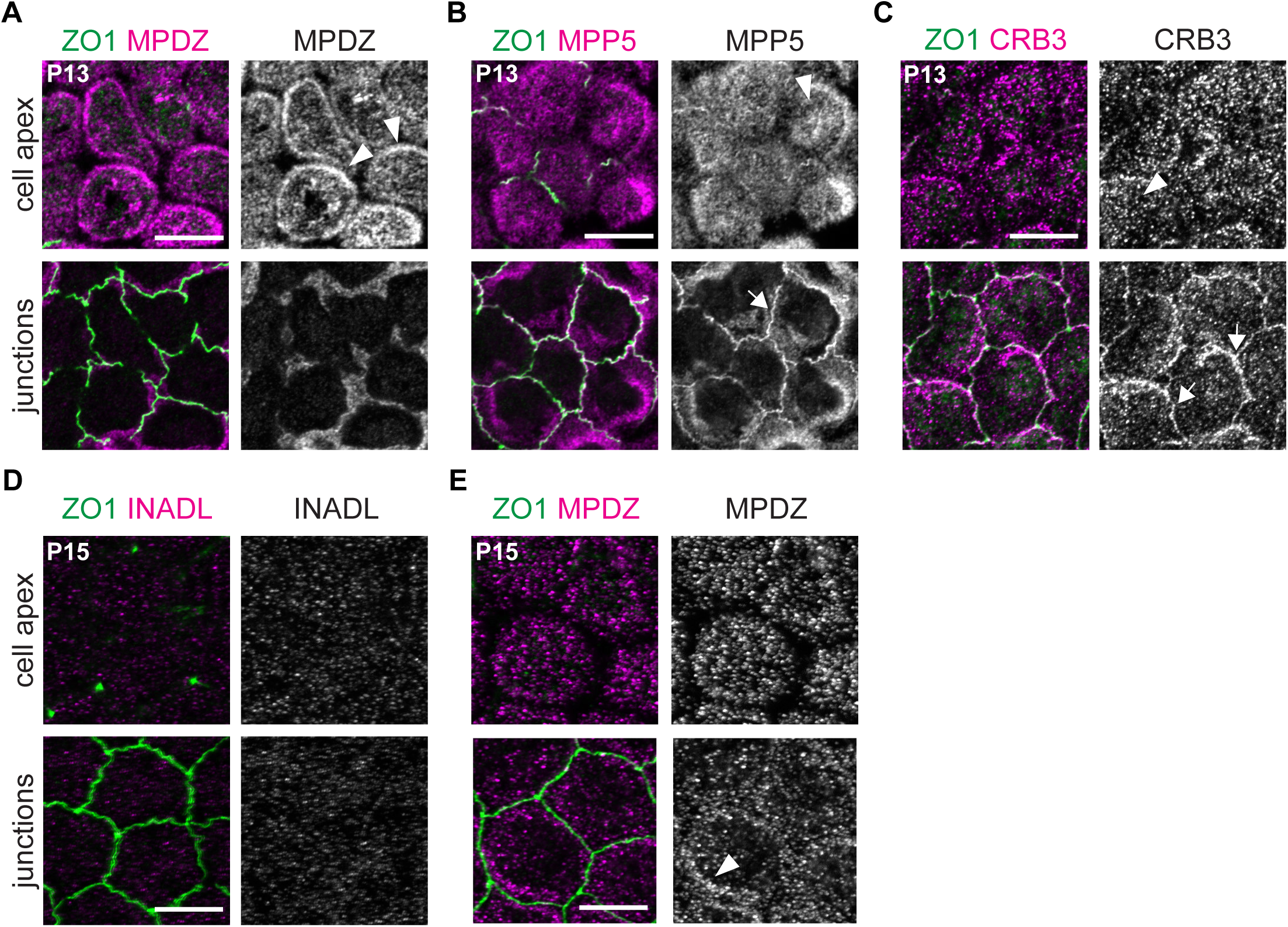
MPDZ, MPP5 and CRB3 localization in the epithelium of the hindbrain choroid plexus. **A-E)** Immunolabeling in P13-P15 wild-type 4^th^ ventricle choroid plexus (projection of a confocal stack) where ZO1 defines the apical junctions. A projection of the most apical Z slices in the stack (1 μm depth) is used to visualize the apical membrane (top panels, “cell apex”). A projection of adjacent, more basal Z slices in the same confocal stack (1 μm depth) is used to visualize the apical junctions (bottom panels, “junctions”). **A)** MPDZ is enriched at the apical membrane where it accumulates near the junctions (arrowheads), but MPDZ is not itself co-localized with ZO1 at apical junctions. **B-C)** In contrast, MPP5 (B) and CRB3 (C) occupy the apical membrane like MPDZ (arrowheads), but also colocalize with ZO1 at the apical junctions (arrows). **D)** INADL is not detected either at the apical membrane or at apical junctions in the choroid plexus epithelium. **E)** An independent experiment at P15 confirmed that MPDZ is enriched at the apical membrane where it accumulates near the junctions (arrowheads), but MPDZ is not itself co-localized with ZO1 at apical junctions. Scale bars are 10μm.

**Supp. Table 1.**
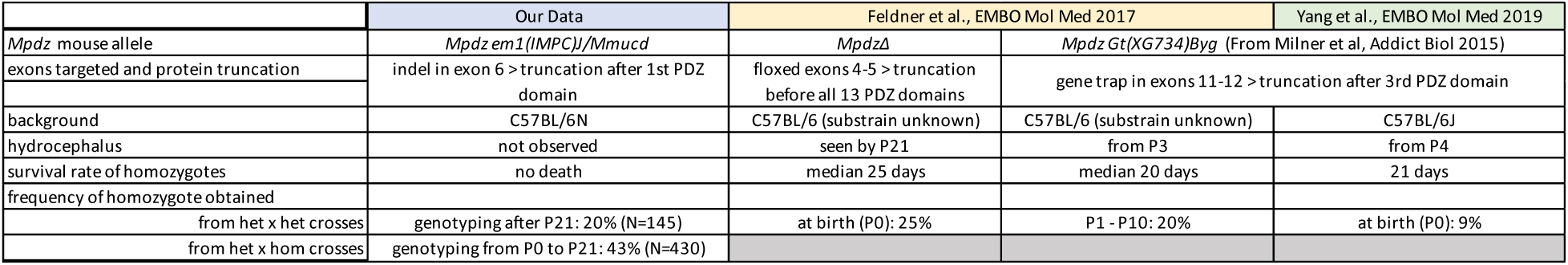

## Notes

### Competing Interest Statement

The authors have declared no competing interest.

### Summary of Updates

We lowered the figure panels so that the biorxiv-added header does not overlap

